# A cucumber green mottle mosaic virus vector for virus-induced gene silencing in cucurbit plants

**DOI:** 10.1101/813741

**Authors:** Mei Liu, Zhiling Liang, Miguel A. Aranda, Ni Hong, Liming Liu, Baoshan Kang, Qinsheng Gu

**Affiliations:** Zhengzhou Fruit Research Institute, Chinese Academy of Agricultural Sciences, Zhengzhou 450009, P.R. China; Huazhong Agricultural University, Wuhan 430070, P.R. China; College of Plant Health and Medicine, The Key Laboratory of Integrated Crop Pest Management of Shandong Province, Qingdao Agricultural University, Qingdao, 266109, China; Centro de Edafologia y Biologia Aplicada del Segura (CEBAS)-CSIC, Apdo. Correos 164, 30100 Espinardo, Murcia, Spain

**Author notes:** **Author contributions:** Qinsheng Gu and Mei Liu conceived the original idea and designed the experiments. Mei Liu, Zhiling Lian, Liming Liu and Baoshan Kang performed the experiments and analyzed the data. Mei Liu wrote the manuscript. Qinsheng Gu, Ni Hong and Miguel A. Aranda revised the manuscript. All authors have read and approved the manuscript. Qinsheng Gu agrees to serve as the author responsible for contact and ensures communication.

**Keywords:** Cucumber green mottle mosaic virus, viral vector, virus-induced gene silencing, cucurbit plants

## Abstract

Cucurbits produce fruits or vegetables that have great dietary importance and economic significance worldwide. The published genomes of at least 11 cucurbit species are boosting gene mining and novel breeding strategies, however genetic transformation in cucurbits is impractical as a tool for gene function validation due to low transformation efficiencies. Virus-induced gene silencing (VIGS) is a potential alternative tool. So far, very few ideal VIGS vectors are available for cucurbits. Here, we describe a new VIGS vector derived from cucumber green mottle mosaic virus (CGMMV), a monopartite virus that infects cucurbits naturally. We show that the CGMMV vector is competent to induce efficient silencing of the phytoene desaturase (*PDS*) gene in the model plant *Nicotiana benthamiana* and in cucurbits, including watermelon, melon, cucumber and bottle gourd. Infection with the CGMMV vector harboring *PDS* sequences of 69-300 bp in length in the form of sense-oriented or hairpin cDNAs resulted in photobleaching phenotypes in *N. benthamiana* and cucurbits by *PDS* silencing. Additional results reflect that silencing of the *PDS* gene could persist for over two months and the silencing effect of CGMMV-based vectors could be passaged. These results demonstrate that CGMMV vector could serve as a powerful and easy-to-use tool for characterizing gene function in cucurbits.

**One sentence summary:** A CGMMV-based vector enables gene function studies in cucurbits, an extremely low efficiency species for genetic transformation.

## Introduction

The family *Cucurbitaceae* is second only after the *Solanaceae* for its economic importance among horticultural species worldwide, containing about 1000 species in 96 genera (Renner and Schaefer, 2016). Cucurbits are generally prized for their delicious fruits, which might be low in nutritional value, but can be significant dietary sources of minerals and vitamins, some even with medical values. Watermelon (*Citrullus lanatus*), melon (*Cucumis melo*), cucumber (*Cucumis sativus*) and bottle gourd (*Lagenaria siceraria*) all belong to the family *Cucurbitaceae* with a significant impact on human nutrition (Grumet et al., 2017).

With the increase of consumer’s demand for high-quality fruits and vegetables and the improvement of agricultural production, it is urgent to explore genes encoding important agronomic traits in crop species, in order to breed elite, disease-resistant and featured varieties. So far, 11 reference genomes of cucurbit species (Zheng et al., 2019) including watermelon (Guo et al., 2013), melon (Garcia-Mas et al., 2012) and cucumber (Huang et al., 2009) have been published, which have boosted gene mining and gene function research. However, the genetic transformation of cucurbit plants is time-consuming and labor-intensive, with extremely low efficiencies (Choi et al., 1994). As a tool for rapid gene function validation, virus-induced gene silencing (VIGS) is a good alternative to gene transformation because of its simplicity, high efficiency, and high throughput.

Gene silencing comprises transcriptional gene silencing (TGS) and post-transcriptional gene silencing (PTGS). VIGS, a type of PTGS, is a natural defense reaction that exists in a broad range of organisms. It confers resistance to foreign nucleic acid invasion through PTGS at the RNA level. Because it can silence a specific gene, leading to the loss of function of this gene, the potential of VIGS as a tool to analyze gene function has been quickly recognized (Baulcombe, 1999).

In the past decades, a large number of viral vectors had been developed as powerful tools for the functional verification of genes in plants (Ruiz et al., 1998; Liu et al., 2002; Ding et al., 2006; Igarashi et al., 2009; Zhang et al., 2010; Sempere et al., 2011; Liu et al., 2016; Wang et al., 2016). To date, three different RNA viruses have been developed as vectors for VIGS in cucurbit species, including apple latent spherical virus (ALSV) (Igarashi et al., 2009), tobacco ringspot virus (TRSV) (Zhao et al., 2016) and tobacco rattle virus (TRV) (Bu et al., 2019; Liao et al., 2019). However, very few applications of these vectors have been reported, implying that they have not been widely adopted for cucurbit gene function analyses. This might be related to their limited host range among cucurbits, cumbersome inoculation approaches and/or short silencing periods associated with insert instability. As a result, it is urgent to develop a vector with a wider range of cucurbit hosts, ease of inoculation, high silencing efficiency and long-lasting gene silencing in cucurbit plants.

Cucumber green mottle mosaic virus (CGMMV) is an important pathogen infecting cucurbit plants in natural conditions (Dombrovsky et al., 2017). We have successfully constructed a full-length infectious clone of CGMMV, which can systemically infect plants of various cucurbit species such as watermelon, melon, cucumber and bottle gourd (Liu et al., 2017), making it a good candidate for VIGS vector development in cucurbits. CGMMV is a member of the genus *Tobamovirus*, and has a positive single-stranded genomic RNA of approximately 6.4 kb (Ugaki et al., 1991). The CGMMV genome possesses four open reading frames (ORFs) encoding two replication-related proteins, one movement protein (MP), and one coat protein (CP). Only the 129 KDa and 186 KDa of replication-related proteins are translated directly from the genomic RNA, whereas the 29-KDa MP and the 17.4-KDa CP are translated from two subgenomic RNAs. There is an overlap between the MP and CP ORFs (Ugaki et al., 1991). Viral vectors based on CGMMV for expressing foreign genes have been constructed. Multiple cloning sites (MCS) were inserted adjacent to the CP ORF, and the CP stop codon was altered to express the hepatitis B surface antigen and a Dengue virus Epitope so that 20 and 44 foreign amino acids, respectively, were expressed (Ooi et al., 2006; Teoh et al., 2009). Tobacco mosaic virus (TMV), another member of the genus *Tobamovirus*, has been widely studied as a model in this genus. TMV has successfully been developed as a VIGS vector by including an additional duplicated copy of the CP subgenomic promoter (SGP) in the viral genome (Kumagai et al., 1995).The CGMMV genome is similar to that of TMV, and thus it was thought that methods similar to those used for TMV could be used to create vectors based on CGMMV; unfortunately, results in this regard varied largely (Zheng et al., 2015; Jailani et al., 2017), therefore the strategy of SGP duplication and information on the subgenomic promoter have not been fully exploited for constructing CGMMV-based viral vectors.

CGMMV has not been reported for its development for VIGS, although it has been exploited as a transient gene expression vector by readthrough translation or adding an additional subgenomic CP promoter. In this study, we developed a new CGMMV-based VIGS vector, which produces very mild viral symptoms and efficiently triggers gene silencing in the model plant *N. benthamiana* and cucurbit plants such as watermelon, melon, cucumber and bottle gourd.

## Materials and methods

### Plant materials

The CGMMV experimental host *N. benthamiana* and cucurbits hosts (watermelon, melon, cucumber and bottle gourd) were used for VIGS of the *PDS* gene by CGMMV vectors in this study. Watermelon (Zhengkang 2), melon (Baimei), cucumber (Jinyan 4) and bottle gourd (Yongzhen1) seeds were obtained from Zhengzhou Fruit Research Institute (Zhengzhou, China), Xinjiang Academy of Agricultural Sciences (Xinjiang, China), Tianjin Academy of Agricultural Sciences (Tianjin, China) and Ningbo Academy of Agricultural Sciences (Ningbo, China), respectively. All cucurbit seeds were soaked in sterile water for 3 ∼ 4 hours at 50 °C, then placed in Petri plates containing wetted filter cotton gauze at 28 °C in darkness until seeds were germinated. Germinated seeds were planted into pots with nutrient matrix and grown in a growth chamber under 16 h light at 28°C / 8 h dark at approximately 22°C. The same conditions were used to grow inoculated plants (see below) with CGMMV vectors.

### Construction of the CGMMV-based vectors

pV1a23 was constructed by site-directed mutagenic PCR. A DNA fragment of about 7000 bp, consisting of the 5’ end of the pXT1-CGMMV, was amplified using pXT1-CGMMV as a template with primer pairs Del*Hind*III-X/ S159Z-S (Table S1); Another DNA fragment of about 4000 bp, consisting of the 3’ end of the pXT1-CGMMV, was amplified using pXT1-CGMMV as a template with primer pairs S159Z-X/Del*Hind*III-S (Table S1). These two fragments were ligated by homologous recombination. The resulting construct was named as pV1a23.

PCR was performed with primers CP-TC-F and CP-TC-R to remove the CP start codon of the pXT1-CGMMV (Table S1), resulting in the single-nucleotide substitution ATG to ACG. The resulting construct was named pXT1-CGACG. pV61, pV92, pV112 and pV190 VIGS vectors were constructed by site-directed mutagenic PCR as well. DNA fragment 1 containing CGMMV nt 1- (5711∼5840) (GenBank accession: KY753929) was amplified using pXT1-CGMMV as a template with primer pairs PXT1-F / (27B-34-R, 58B-34-R, 78B-34-R or 78B-99-R), whereas DNA fragment 2 containing CGMMV nt 5651/5716 – 6423 was amplified using pXT1-CGACG as a template with primer pairs PXT1-R /(27B-34-F, 58B-34-F, 78B-34-F or 78B-99-F) (Table S1). These two fragments were ligated by homologous recombination. The resulting vectors pV61, pV92, pV112 and pV190 are pXT1-CGMMV derivatives that include a duplicated copy of 61-bp, 92-bp, 112-bp and 190-bp, respectively, putative CGMMV CP SGP and a single restriction site (*Bam*HI) between the duplicated CP SGP.

### Insertion of different *PDS* fragments into the CGMMV-based vector

For a VIGS test with *PDS* as the target gene, 114-, 213-, and 300-bp cucurbit *PDS* fragments were inserted into digested pV1a23 with *Hind*III in sense orientation to produce *PDS* silencing constructs pV1a23-PDS11414, pV1a23-PDS21313 and pV1a23-PDS30000, respectively. Three primer sets CuPDS-*Hind*III-F/R, CuPDS-*Hind*III-2F/ 2R and CuPDS-*Hind*III-3F/3R were designed to amplify 114-, 213- and 300-bp fragments of the cucurbit *PDS* gene, respectively (Table S1). Similarly, a series of pV92, pV112 and pV190-based vectors harboring different *PDS* fragments of varied sizes were constructed. Seven primer sets 58-150-F/R, 78-34-150F/R, 78-150-F/R, 58-213-F/-R, 78-34-213F/R, 78-213-F/R, 78-300-F/R were used for amplifying 150-, 213- and 300-bp fragments of the cucurbit *PDS* gene, respectively (Table S1). Two primer sets 78-146N-F/R and 78-215N-F/R were used for amplifying 146- and 215-bp fragments of the *N. benthamiana PDS* gene, respectively (Table S1). The resulting ten pV92, pV112 and pV190-derived constructs were named pV92-PDS150, pV92-PDS213, pV112-PDS150, pV112-PDS213, pV190-PDS150, pV190-PDS213, pV190-PDS300, pV190-NbPDS146 and pV190-NbPDS215.

pV190-PDS69, a construct carrying a 69-bp fragment (dsRNA hairpin structure) of the cucurbit *PDS* gene, was constructed using three primer sets 78-69P-X/78-69P-S, CG-4R/CG-4F, 3R/TxR∼R (Table S1). DNA fragment 1 of 870 bp, DNA fragment 2 of 1438 bp and DNA fragment 3 of 9 kb, were PCR-amplified using pV190 as template and primers 78-69P-X/CG-4R, CG-4F/78-69P-S and 3R/ TxR∼R, respectively. DNA fragment 1, DNA fragment 2 and DNA fragment 3 were ligated by homologous recombination.

### Agroinfiltration and sap inoculation

All constructs were introduced into *Agrobacterium tumefaciens* strain GV3101 by freeze-thaw transformation, then single clones were picked up and transferred into 200 µL LB liquid media containing kanamycin (50 µg mL^−1^) and rifampicin (50 µg mL^−1^) and cultured overnight in a shaker at 28℃. The bacterium culture was mixed with LB at a 1:100 ratio and cultured in a shaker overnight, followed by centrifugation at 6000×*g* for 5 min to collect the bacteria. The bacteria were resuspended in inducing buffer solution containing 10 mmol L^−1^ MgCl_2_, 10 mmol L^−1^ MES, and 100 µmol L^−1^ Acetosyringone, and the final OD_600_ value was adjusted to 0.8 ∼ 1. The cells were maintained at room temperature (25℃) for at least 2 h before agroinoculation. The upper 2∼3 leaves of *N. benthamiana* at the 6∼8 leaf stage and cotyledons from 14-day-old cucurbit seedlings were infiltrated with the *A. tumefaciens* suspension using a 1-mL syringe.

In order to verify whether the silencing effect of these vectors could be passaged, the sap from leaves of the agroinfiltrated melon plants displaying obvious photobleaching was used to rub-inoculate cotyledons and the first true leaf (L1) of the melon plants. Each experiment was repeated at least three times, with 9 plants for each construct in each experiment.

### DAS-ELISA and RT-PCR

After agroinfiltration and sap inoculation, CGMMV in inoculated plants was detected by DAS-ELISA and RT-PCR at specific time points. DAS-ELISA was performed to detect CGMMV accumulation using an ELISA kit (Adgen, Auchincruive, UK). For RT-PCR, total RNA was extracted from cucurbit (watermelon, melon, cucumber and bottle gourd) and *N. benthamiana* leaf tissues using the RNA simple kit (Tiangen Biotech, Beijing, China) and then first-strand cDNA was synthesized from 1 μg total RNA using an oligo dT primer according to the protocol of PrimescriptII RT (TAKARA). PCR was performed with primer set 5574F and 3UTR that flanked the foreign insert to detect CGMMV and asses the stability of the pV190 and foreign inserts of CGMMV-based vectors (Table S1).

### qRT-PCR analysis

qRT-PCR was performed to measure the mRNA expression level of the endogenous *PDS* genes using the SYBR Green I Master (Roche) in either *N. benthamiana* or cucurbit plants inoculated with CGMMV-based vectors at specific time points. The first-strand cDNA was synthesized from 1μg total RNA using an oligo dT primer according to the protocol of PrimeScript™RT reagent Kit with gDNA Eraser (TAKARA). The expression level of *PDS* of cucumber, melon, gourd and watermelon was determined using primer sets CuPDS-679F/CuPDS-906R and wate-q-F/R, respectively, designed to prime outside the region targeted for silencing (Table S1). Expression of the actin gene by primer set cumsactin-F/R (Table S1) was used as an internal control of cucumber, melon, gourd plants. The *ClCAC* gene was used as an internal control of watermelon plants using primer pairs Cla016178-F/R (Kong et al., 2015). The primer set NbPDS-qF/R was designed for detecting the expression of *PDS* in *N. benthamiana*, and the expression of the GAPDH gene analyzed by the primer set GAPDH-qRT-F/R was referred as an internal control (Table S1). The expression *PDS* was calculated using the 2^−ΔΔCT^ method (Livak and Schmittgen, 2001). The expression level of *PDS* in the negative control (pV190) was set to an arbitrary value (1.0) to calculate the relative expression levels of the other samples, with 3 replicates used for each sample.

## Results

### Construction of a set of CGMMV vectors

The selection of the insertion site of the foreign gene fragment is the first step in constructing a VIGS vector. Viable options for CGMMV included placing the insertion site behind the viral MP gene or between the CP gene stop codon and the 3’ non-coding region. For the set of vectors built and tested in this study, we chose to use the latter as a first strategy. Two *Hind*III restriction sites were found in pXT1-CGMMV, one located at the 5’ end and the other at the 3’ end of the CP. We used the latter as the insertion site for constructing our first VIGS vector, pV1a23, which is a pXT1-CGMMV derivative missing the first restriction *Hind*III site and with the 159^th^ amino acid of the CP mutated to a stop codon (Fig. 1A). Cucurbit plants inoculated with this vector showed viral symptoms on upper leaves similar to those of plants inoculated with the pXT1-CGMMV, and CGMMV could be detected by DAS-ELISA and RT-PCR in these leaves (Fig. 1B, C; Table S2).

**Figure 1:**
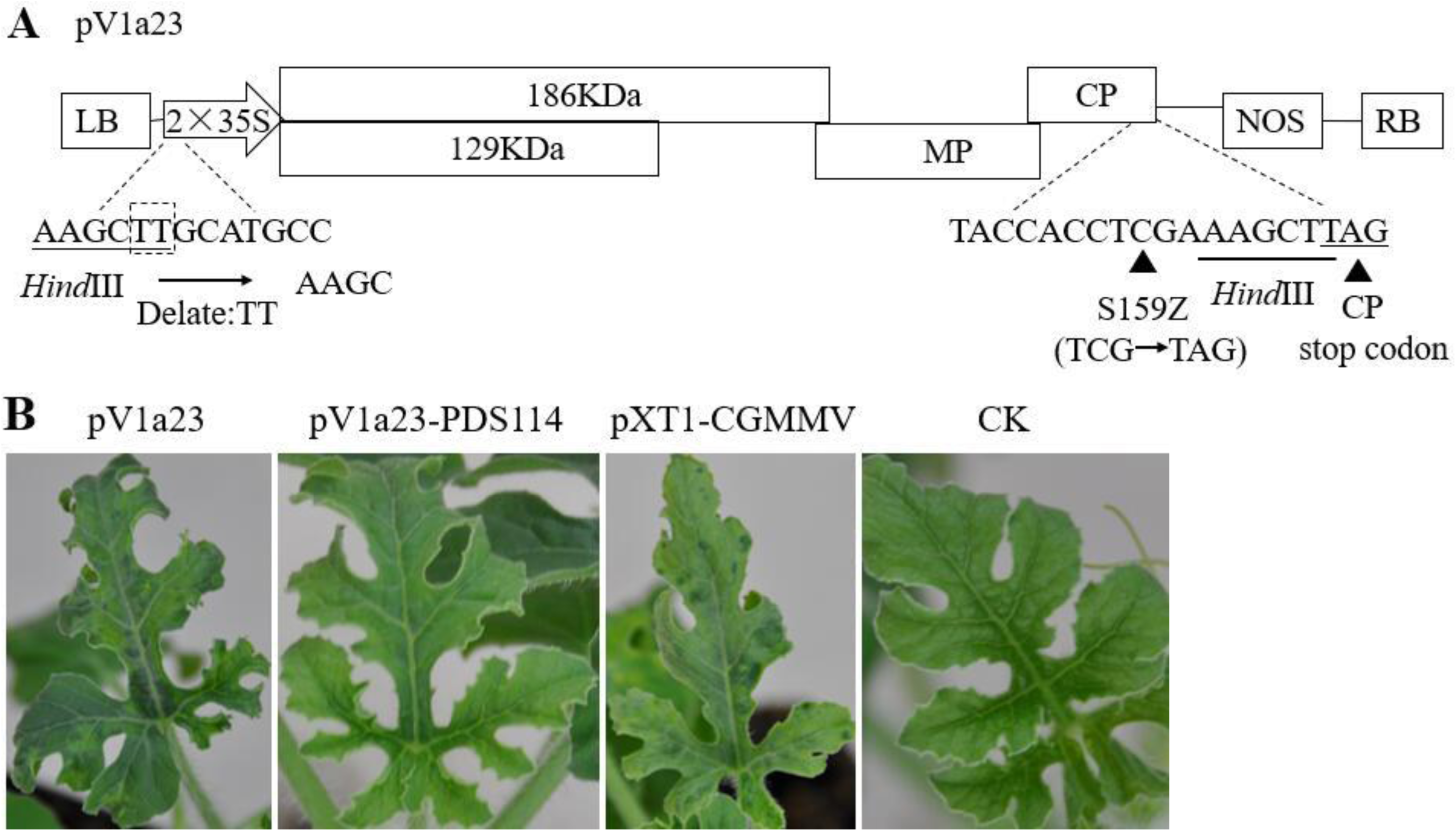
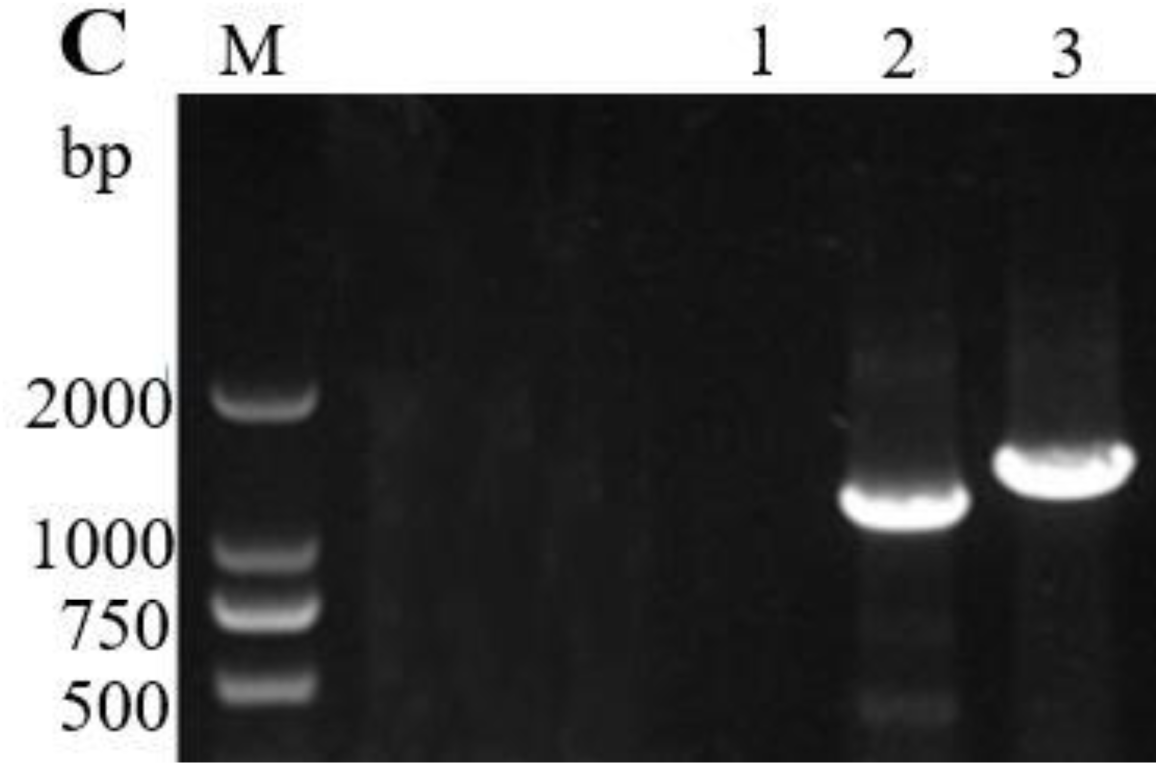
Engineering of CGMMV as a VIGS vector with an insertion site behind the CP. A, Schematic representation of the pV1a23 vector with a restriction enzyme site (*Hind*III) for insertion of gene fragments. B, Viral symptoms on upper non-inoculated leaves caused by pV1a23 similar to those of plants inoculated with the pXT1-CGMMV. Photobleaching was absent on plants inoculated with pV1a23-PDS114. C, RT-PCR detection of viral RNA from pV1a23 and pV1a23-PDS114 in watermelon. M: Marker2000; CK: negative control; 1, 2 and 3 indicate healthy control and plants inoculated with pV1a23 and pV1a23-PDS114, respectively.

CGMMV, similarly to TMV, belongs to the genus *Tobamovirus*, and TMV has been successfully used in VIGS. Thus, we used a second strategy, similar to that used for TMV by including a duplicated copy of the CP SGP in the viral genome (Kumagai et al., 1995) to build vectors pV61, pV92 and pV112; these vectors are pXT1-CGMMV derivatives with different lengths of the CP promoter (61, 92 and 112 nucleotides) and a single restriction site *(Bam*HI) between the duplicated CP promoter (Fig. 2A). Viral symptoms could be observed on upper leaves of plants inoculated with vectors pV92 and pV112, whereas plants inoculated with pV61 did not develop viral symptoms and CGMMV could not be detected by DAS-ELISA (Table S2). These results revealed that vectors pV92 and pV112 have the ability to infect plants systemically while pV61 has not.

**Figure 2:**
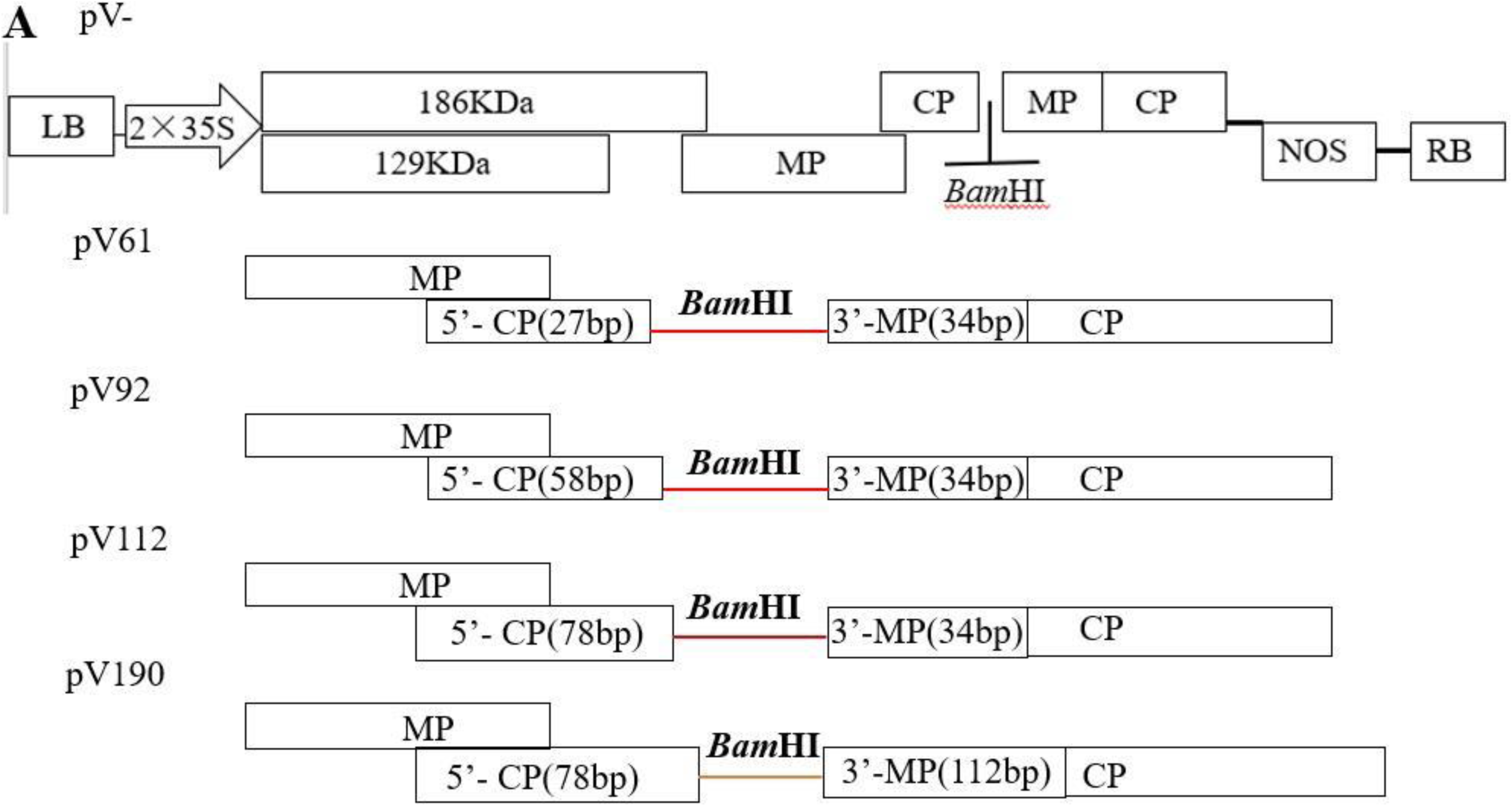
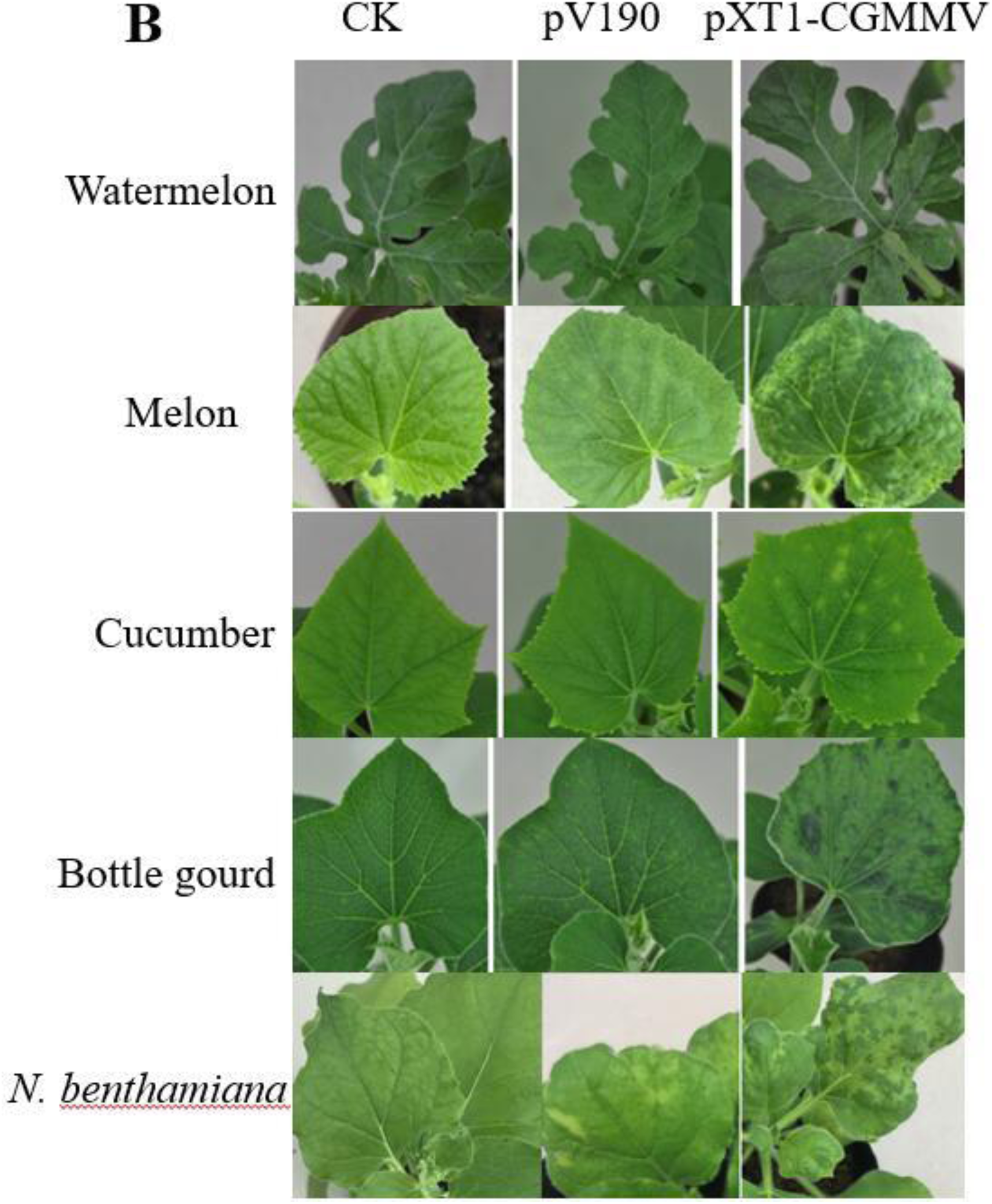
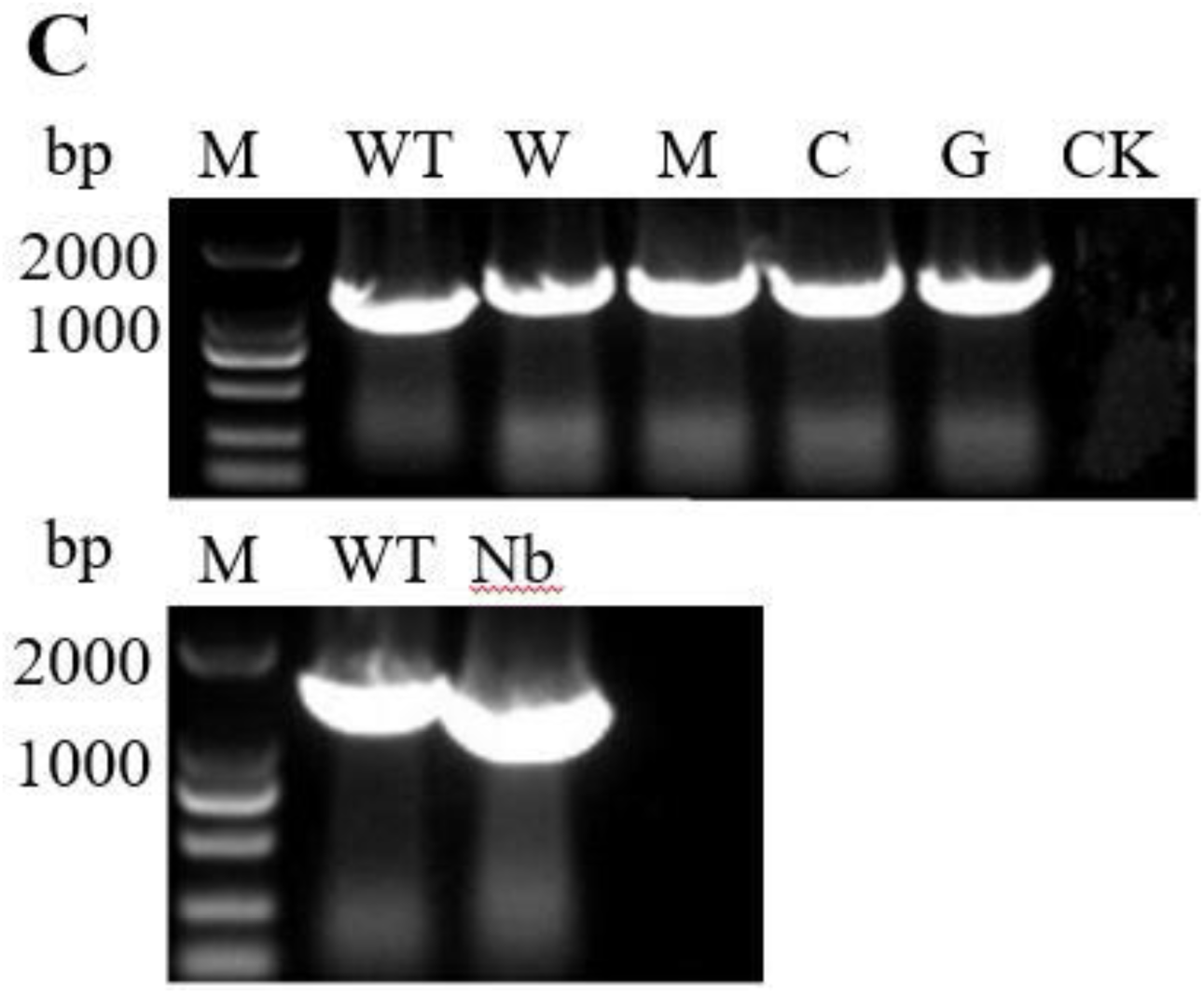
Engineering of CGMMV as a VIGS vector with different size CP subgenomic promoters. A, Schematic representation of pV61, pV92, pV112 and pV190. pV-is a pXT1-CGMMV derivative that contains a direct repeat of the 61-, 92-, 112- and 190-bp putative CGMMV CP subgenomic promoter and a restriction enzyme site (*Bam*HI) between CP subgenomic promoters. B, pV190 caused mild systemic symptoms on cucurbits and *N. benthamiana.* C, RT-PCR detection of viral RNA showing that pXT1-CGMMV and pV190 are infectious in cucurbits and *N. benthamiana*. M: Marker2000; WT: wild type (pXT1-CGMMV); CK: negative control; W, M, C, G and Nb indicate watermelon, melon, cucumber, bottle gourd and *N.benthamiana*, respectively.

Our previous work revealed that the CP RNA transcription level was significantly enhanced when 105 nucleotides were retained before the CP transcription starting site (TSS) and that the sequence from the 71^st^ base to the 91^st^ base upstream of the CP TSS plays a key role in CP SGP activity (Liu et al., 2019). Based on these results, we built pV190, which is a pXT1-CGMMV derivative that contains a direct repeat of the 190-bp putative CGMMV CP SGP and a single restriction site (*Bam*HI) between the duplicated CP SGPs (Fig. 2A). *N. benthamiana* and cucurbit plants inoculated with pV190 developed very mild symptoms on upper leaves, which were clearly milder than those of plants inoculated with the pXT1-CGMMV (Fig. 2B). However, the recombinant pV190 genomic RNA could be detected by RT-PCR (Fig. 2C), indicating that it could replicate and move systemically.

### Silencing effects of *PDS* fragments inserted in the sense orientation or conforming a hairpin

To determine whether pV1a23, pV92, pV112 and pV190 can be used to induce gene silencing in cucurbits, we chose to target *PDS* because it can result in striking photo-bleaching when silenced (Holzberg et al., 2002). Based on an alignment of *PDS* gene coding sequences of watermelon, cucumber, melon and bottle gourd, we designed four primer sets selecting the region with the highest conservation to amplify 114-, 150-, 213- and 300-bp fragments of the cucurbit *PDS* genes. The sequences similarity of the 114-, 150- and 213-bp fragments in the four cucurbit species was approximately 97% (Fig. S1A∼C). The sequence similarity of the 300-bp fragments was 98.4%, but the fragment from watermelon contains an insertion of 30 bp (Fig.S1D). The *PDS* fragments of 114-bp, 213-bp and 300-bp were inserted in the sense orientation at the *Hind*III cloning site of pV1a23 to produce pV1a23-PDS114, pV1a23-PDS213 and pV1a23-PDS300, respectively. To verify the silencing efficiency of these vectors, cucurbit plants were subjected to *Agrobacterium*-mediated inoculation. Uninoculated leaves of inoculated plants with pV1a23-PDS213 or pV1a23-PDS300 did not show any symptoms at 14 days post inoculation (dpi) and 21 dpi. Viral symptoms could be clearly observed in systemically infected leaves of all infected plants with pV1a23-PDS114; however, the *PDS*-silencing photobleaching phenotype could not be observed (Fig. 1B). DAS-ELISA showed the presence of CGMMV in uninoculated leaves of inoculated plants with pV1a23-PDS114 and in leaves inoculated with pV1a23-PDS213, however, the presence of CGMMV could not be detected in leaves of plants inoculated with pV1a23-PDS300 (Table S2). We reasoned that the absence of the photobleaching could be due to the deletion of the 114-bp *PDS* gene fragment. However, RT-PCR showed that the 114-bp *PDS* gene fragment was stable (Fig. 1C). These results revealed that pV1a23-PDS300 lost the ability of systemic and local infection, pV1a23-PDS213 was only able to infect locally and pV1a23-PDS114 could produce systemic and local infection, but failed inducing photobleaching.

The *PDS* fragments of 150 bp and 213 bp were inserted in the sense orientation at the *Bam*HI cloning site of pV92 and pV112 to produce pV92-PDS150, pV92-PDS213 and pV112-PDS150 and pV112-PDS213, respectively. Watermelon, cucumber and melon plants were inoculated with these vectors for testing their ability to induce *PDS* silencing. Photobleaching could be observed in the inoculated plants with pV92-PDS150 and pV112-PDS150 (Table S3). However, pV92-PDS213- and pV112-PDS213-infected plants did not display any photobleached phenotype and the presence of CGMMV in the upper leaves was not observed (Table S3).

The *PDS* fragments of 150 bp, 213 bp and 300 bp were also inserted in the sense orientation at the *Bam*HI cloning site of pV190 to produce pV190-PDS150, pV190-PDS213 and pV190-PDS300, respectively. The cotyledons of watermelon, melon, cucumber and bottle gourd seedlings were inoculated with the above vectors. Photo-bleaching was first observed on the 4^th^ true leaves (L4) in watermelon at about 19 dpi, on the 3^rd^ leaves (L3) in melon and bottle gourd at 12 dpi (Fig. 3A), and on the 5^th^ true leaves of cucumber at 28 dpi (data not shown). Further, photobleaching was observed up to 32, 20 and 39 dpi in watermelon, melon and cucumber plants, respectively (Fig. 3B). About 70% of the inoculated plants showed a photobleaching phenotype. Total RNA was extracted from leaves of the plants inoculated with different vectors displaying the most obvious photobleaching (Fig. 4A) and the accumulation of *PDS* transcripts was quantified by qRT-PCR. The results showed that the expression levels of *PDS* had no significant differences between pV190-infected (EV) and noninfected (NI) leaves, demonstrating that pV190 did not significantly affect *PDS* expression (Fig. 4B). The *PDS* mRNA transcript levels in photobleached leaves was reduced by approximately 79%, 81% and 89% in watermelon, 78%, 76% and 81% in melon, 83%, 87% and 89% in bottle gourd, and 82%, 64% and 88% in cucumber infected with pV190-PDS150, pV190-PDS213 and pV190-PDS300, respectively, compared to plants infected with pV190 (Fig. 4B).

**Figure 3:**
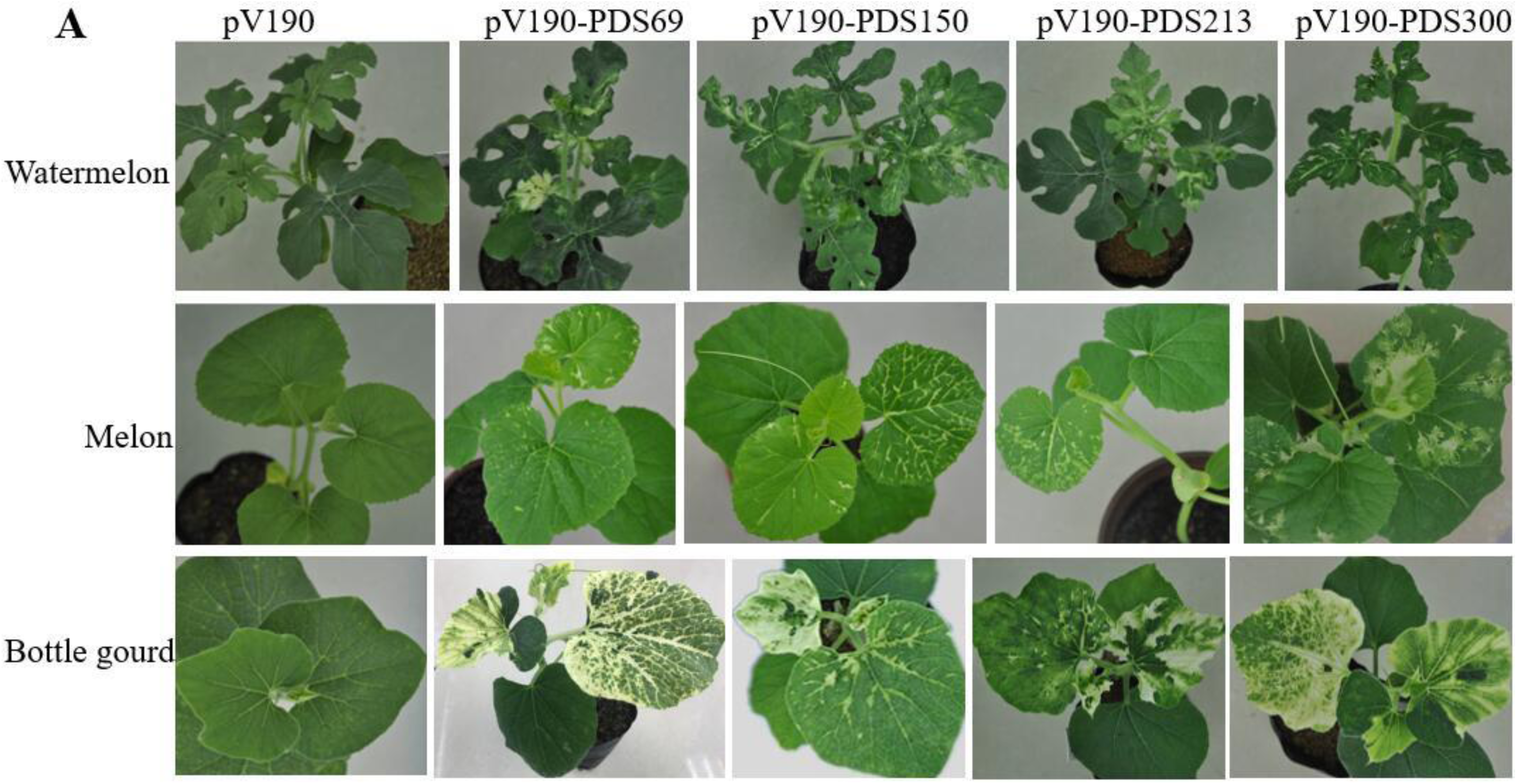
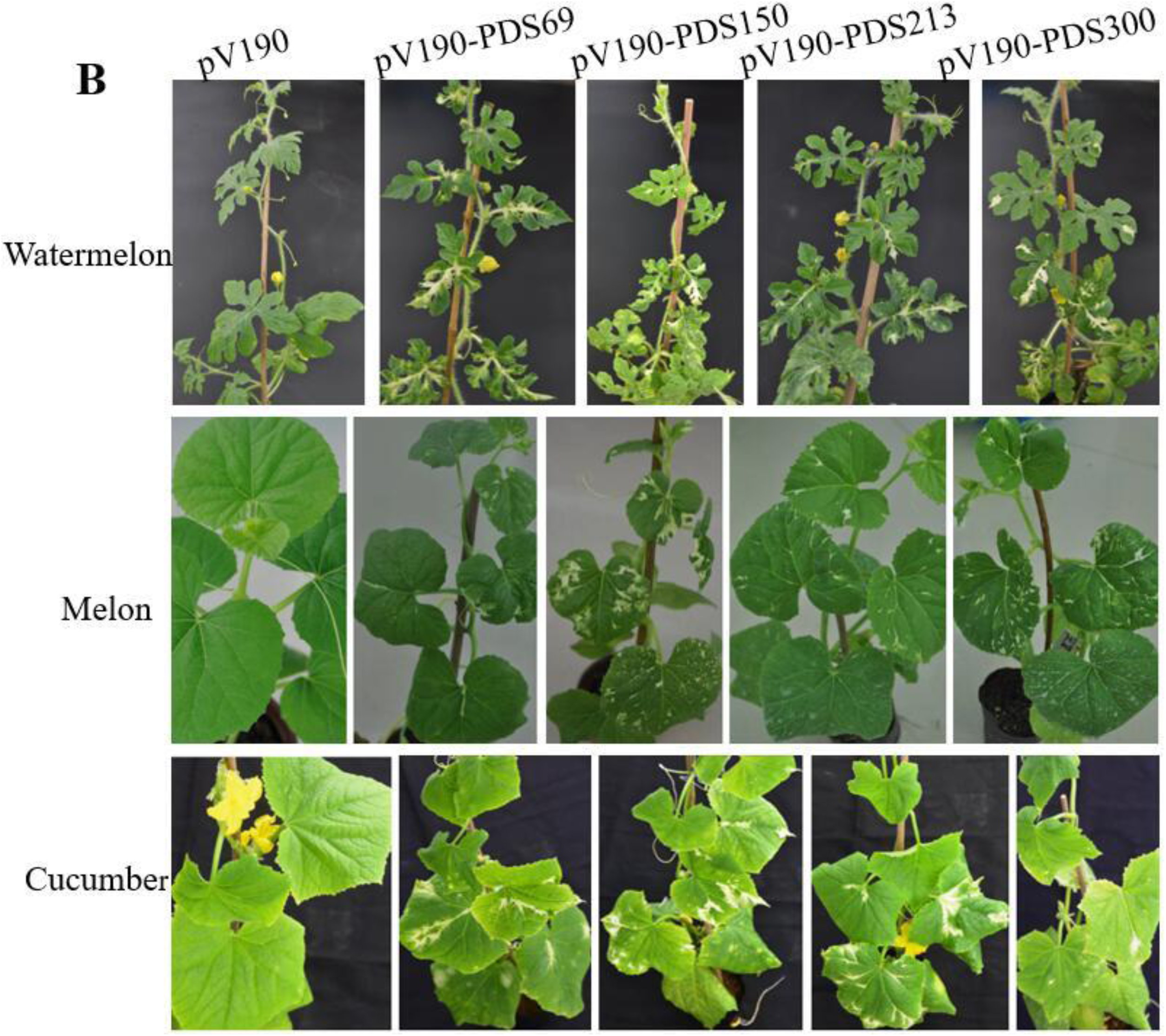
*PDS* silencing using the VIGS vectors pV190-PDS69, pV190-PDS150, pV190-PDS213 and pV190-PDS300. A, Photobleaching was first observed and photographed in watermelon at 19dpi, and in melon and bottle gourd plants at 12dpi. B, Photobleaching was photographed in watermelon at about 32 dpi, in melon at about 20 dpi and in cucumber at about 39 dpi, respectively.

**Figure 4:**
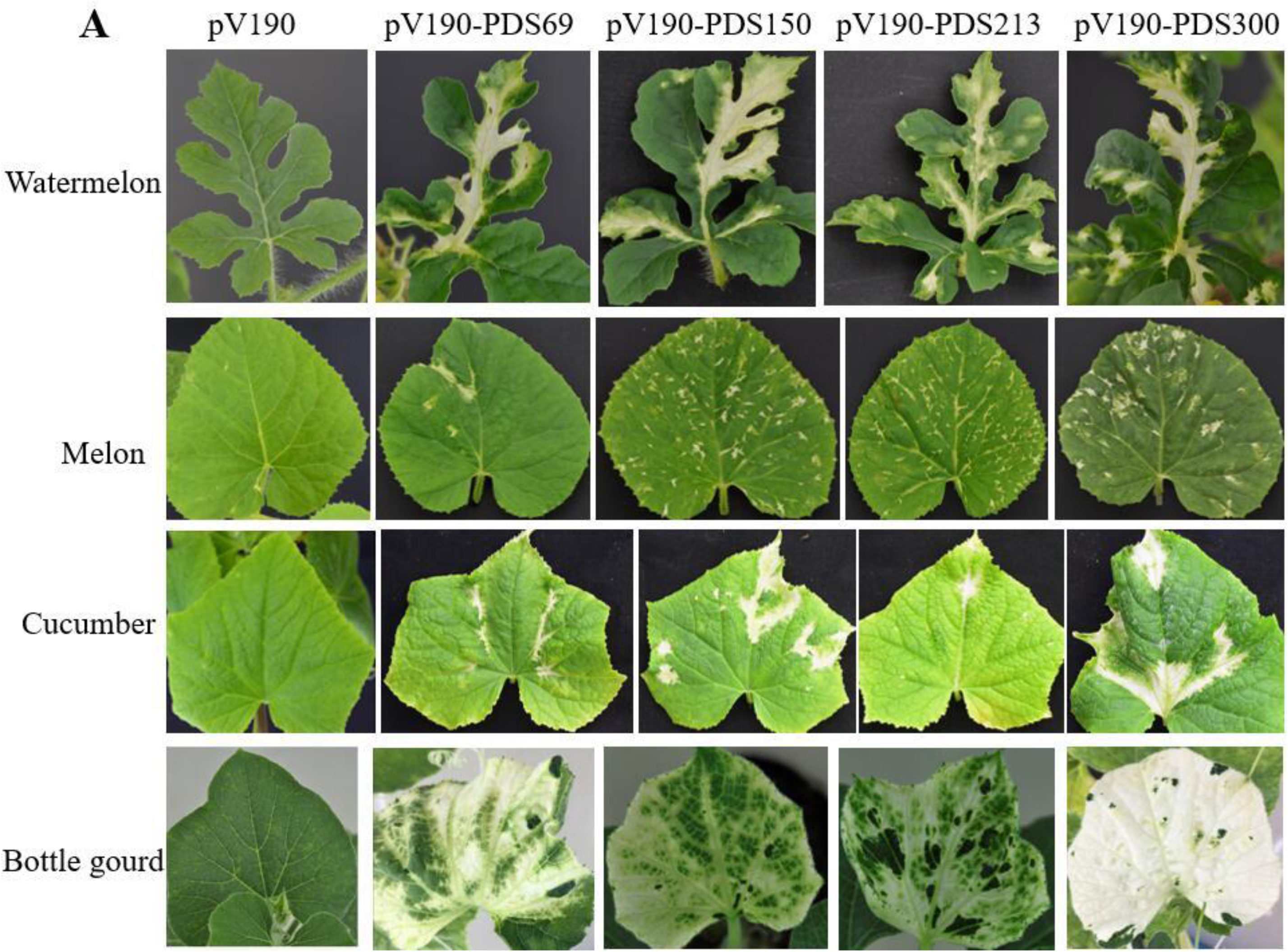
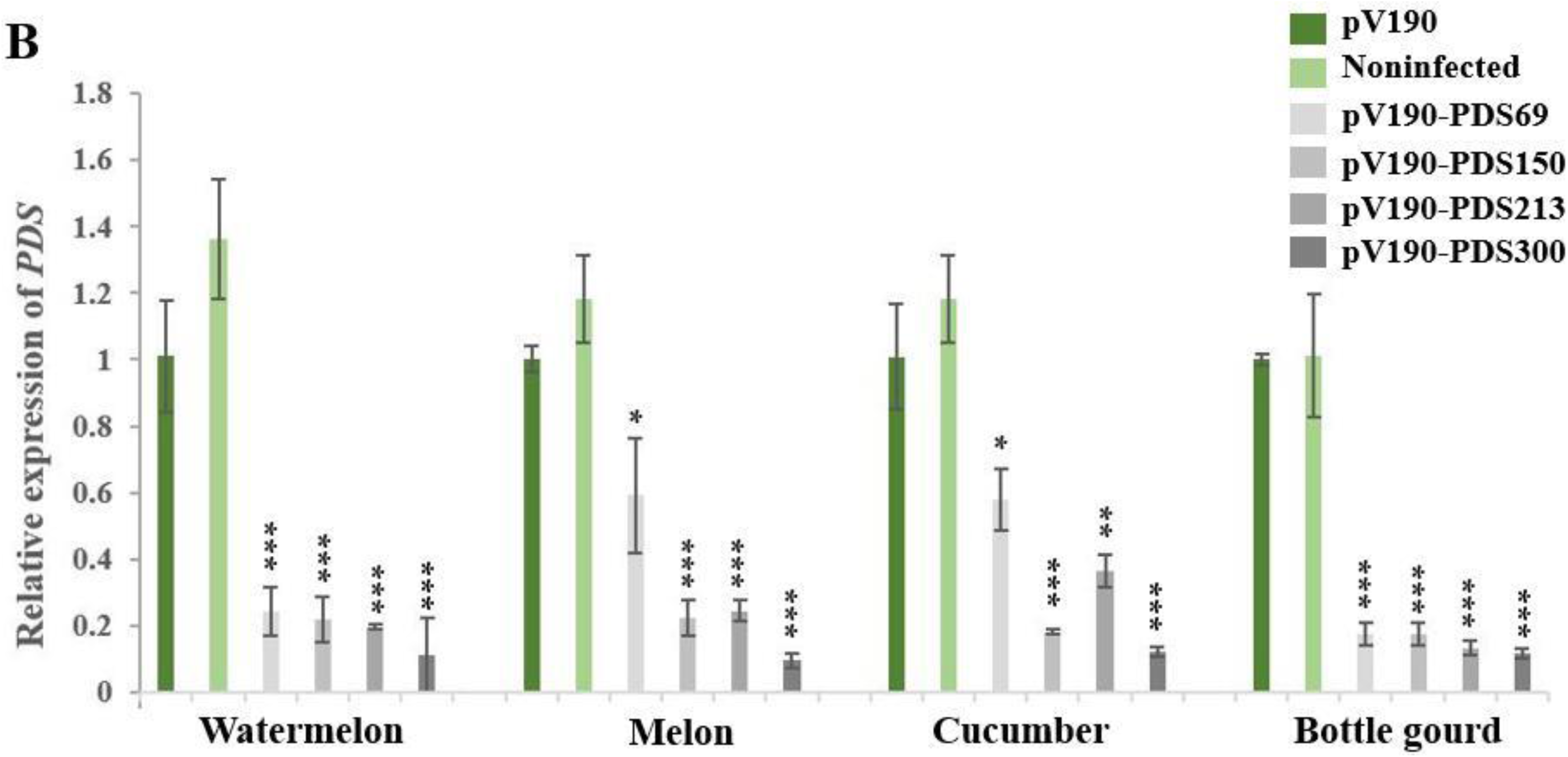
Silencing efficiency of VIGS vectors carrying *PDS* fragments of different sizes on cucurbits. A, indicate the uninoculated leaves displaying most obvious photobleaching on watermelon plants at 32 dpi, on melon at 27 dpi, on cucumber at 39 dpi and on bottle gourd at 34 dpi, respectively. B, Real-time qRT-PCR analysis of *PDS* expression in noninfected (NI), pV190 empty vector (EV), and CGMMV-PDS-infected cucurbit (watermelon, melon, cucumber and bottle gourd) plants. Three technical replicates were performed for each individual sample (*, P < 0.05 and **P < 0.01 ***P < 0.001 compared with the empty vector (pV190) by Student’s t test. Error bars indicate the SD.

To improve silencing efficiency, we inserted a *PDS* fragment forming a hairpin structure of 69-bp into pV190 to produce the pV190-PDS69 vector. We first observed photobleaching on the L5 in watermelon at 17 dpi, on the L2 in melon at 10 dpi and on the L3 in bottle gourd at 11 dpi (Fig. 3). Photobleaching was first observed one or two days earlier in plants infected with pV190-PDS69 than in plants infected with any other vector. The *PDS* mRNA levels declined 76%, 41%, 42% and 83% in watermelon, melon, cucumber and bottle gourd, respectively (Fig. 4B).

### Stability of the 69-300-bp *PDS* fragments in pV190

We observed that the silencing phenotype of the *PDS* gene could persist for over 2 months in bottle gourd (Fig. 5A). Photobleaching was not uniform from bottom to top of bottle gourd leaves (Fig. 5B). To evaluate the stability of the *PDS* fragment in CGMMV-vectors, RT-PCR was performed on total RNA extracted from bottle gourd leaves L6, L7 and L9 for pV190-PDS69, L4 and L11 for pV190-PDS300, L4 and L12 for pV190-PDS213, and L7, L9 and L10 for pV190-PDS300 (Fig. 5B). The result showed the 150-bp and 213-bp *PDS* fragments were stable across all analyzed leaves. The 69-bp dsRNA hairpin structure could not be detected across all leaves, whereas L9 and L10 samples from pV190-PDS300-infected bottle gourd contained deletions of the 300-bp *PDS* fragment to different extents (the deletion in L9 was less than the L10) (Fig. 5C). The relative expression of the *PDS* gene in the above same leaves was measured by qRT-PCR. RT-PCR results corresponded well with the *PDS* relative expression level measured by qRT-PCR, with less silencing observed as the extent of deletions increased (Fig. 5C, D). For instance, pV190-PDS69 caused *PDS* transcripts to be reduced by 83%, 80% and 65% in the L6, L7, L9, respectively, and pV190-PDS300 caused *PDS* transcripts to be reduced by 87%, 81% 73% and 65% in the L7, L8, L9 and L10, respectively (Fig. 5D). Results of stability of the 69-300-bp *PDS* fragments in pV190 in watermelon, melon and cucumber were consistent with those in bottle gourd (data not shown). The expression of *PDS* in the youngest analyzed leaves was still down-regulated (Fig. 5D), indicating that these vectors have sufficient stability to be used to characterize gene functions in cucurbit plants.

**Figure 5:**
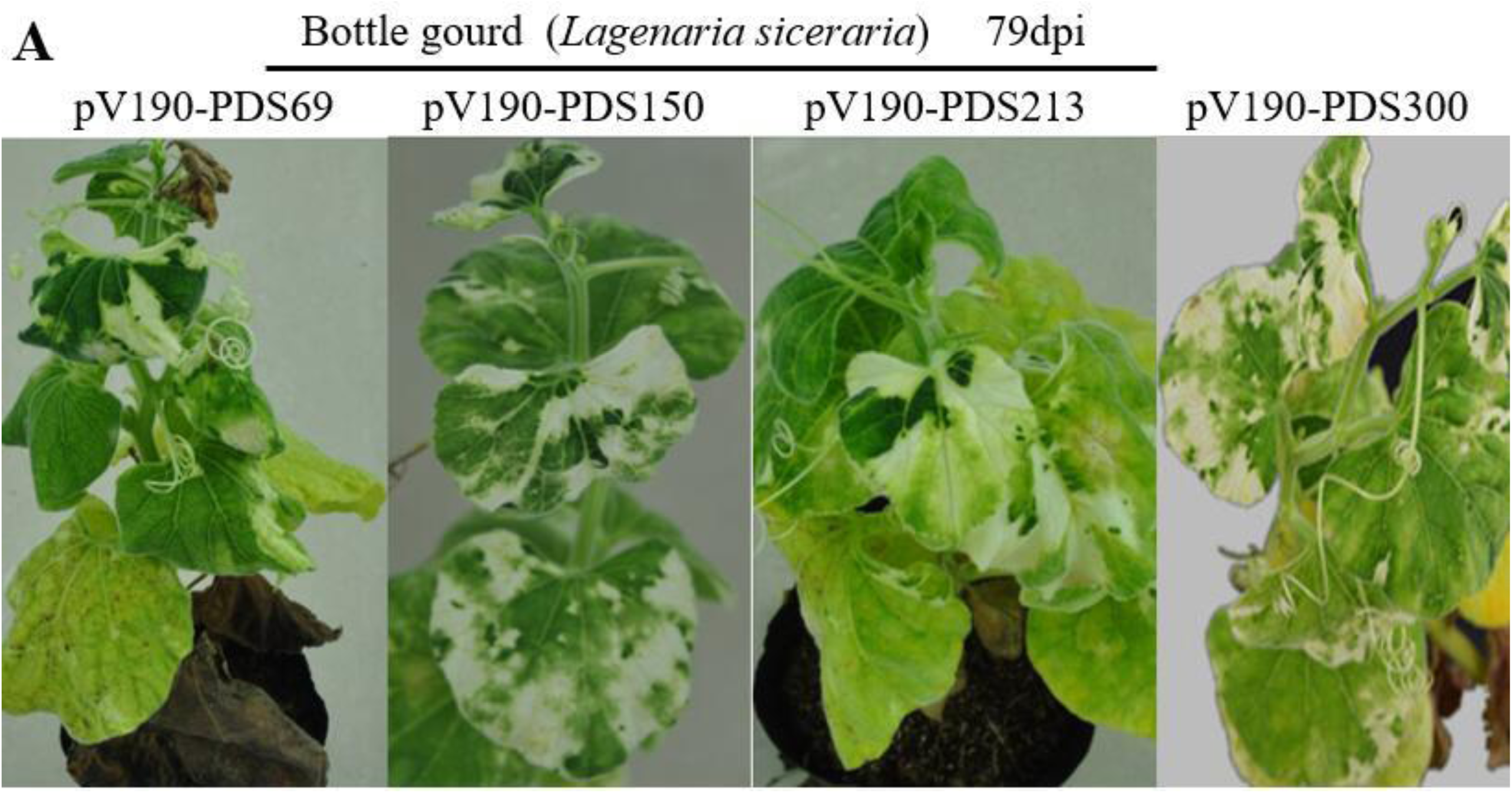
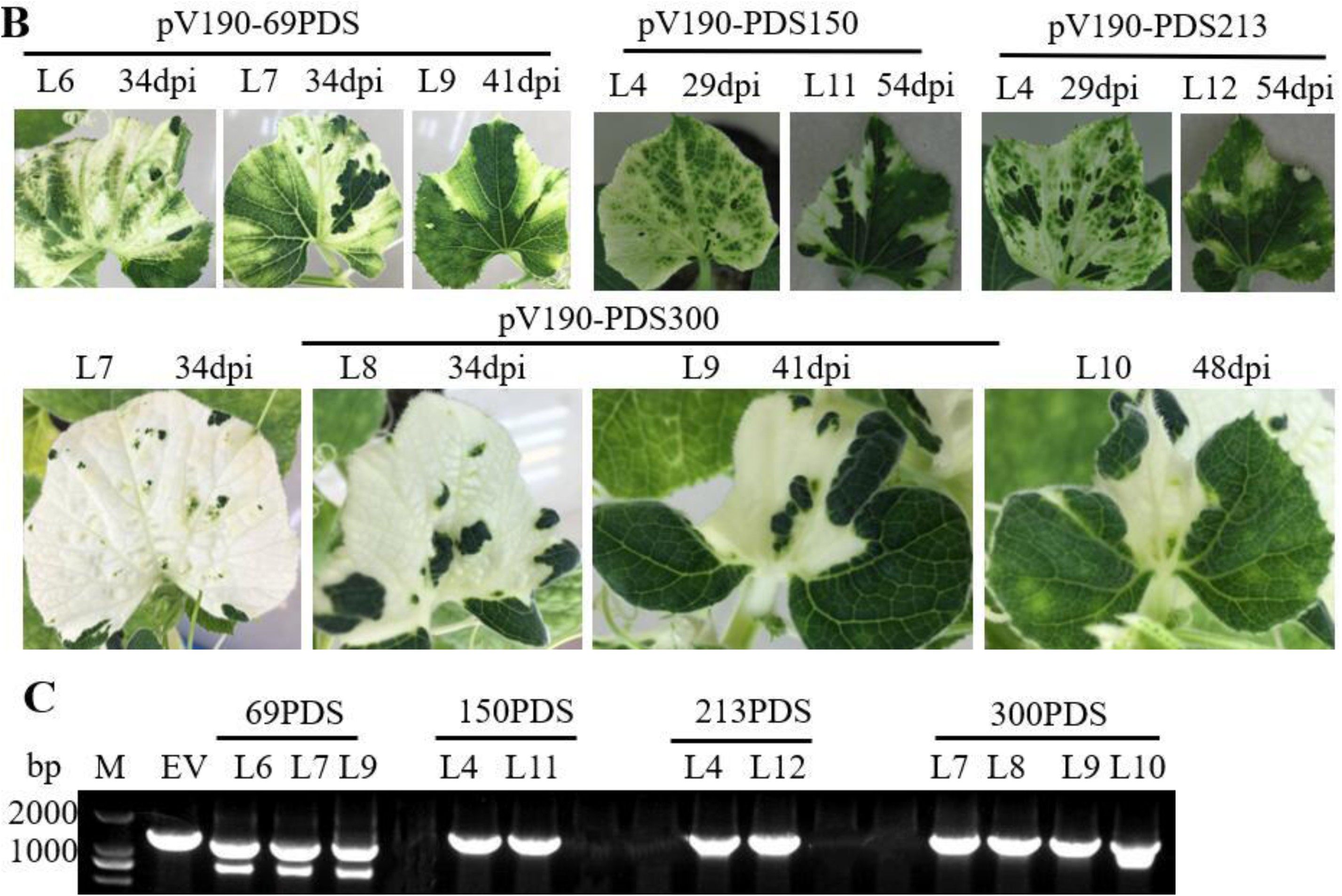
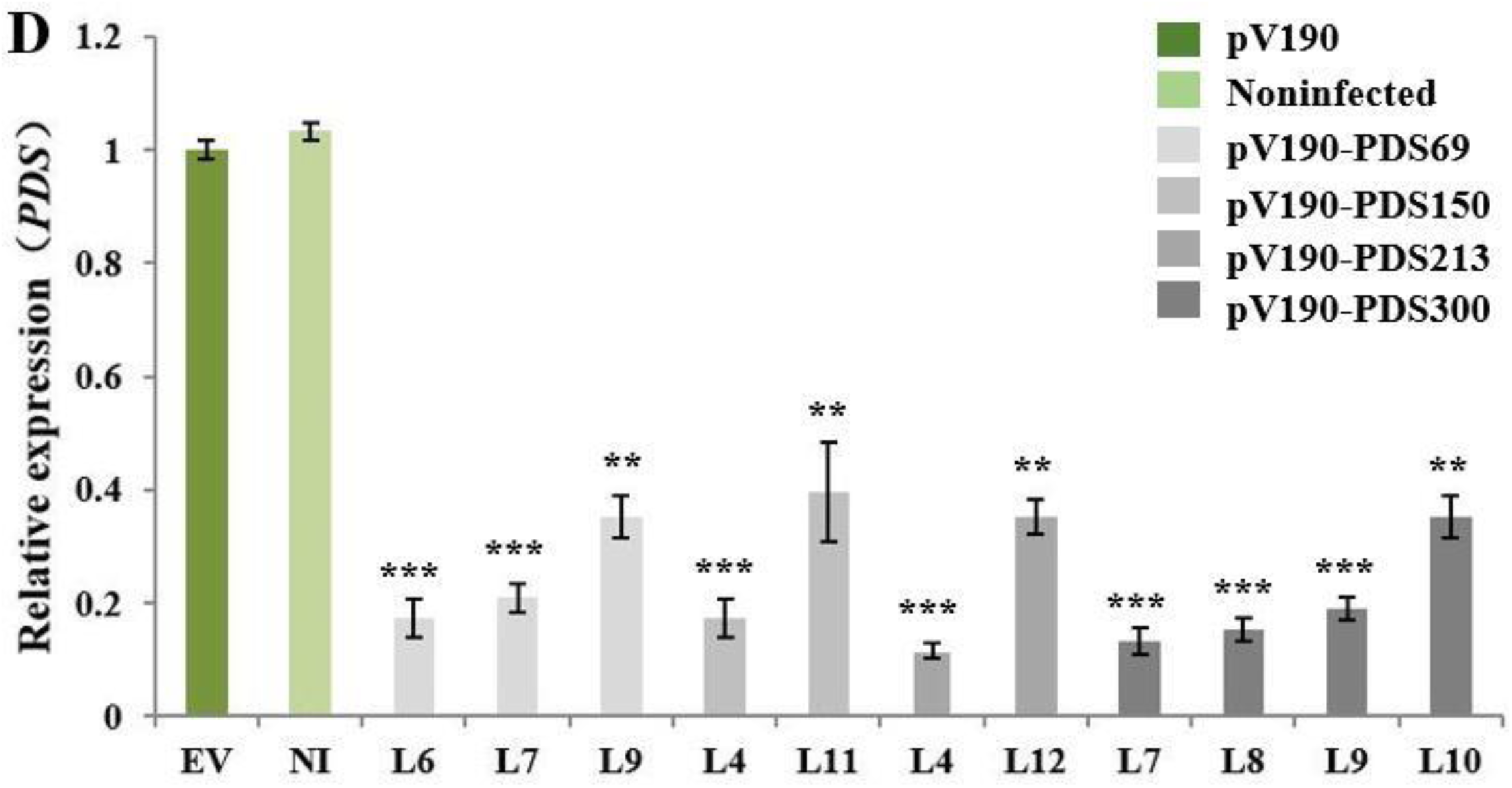
Silencing efficiency and stability of the pV190 VIGS vector with different length inserts in bottle gourd. Fragments of 69 bp (dsRNA hairpin structure), 150 bp, 213 bp, 300 bp were separately cloned into pV190. A, Silencing *PDS* using pV190 on bottle gourd plants produced photobleaching that persisted for over 70 days. B, Photobleaching on newly emerging leaves of bottle gourd plants caused by *PDS* silencing was observed at 29, 34, 41, 48 and 54 dpi, respectively. C, RT-PCR assay to detect the presence of pV190 carrying *PDS* fragments of different sizes in systemic leaves. Samples from the 4^rd^ leaf above the inoculated (L4) were collected at 29 dpi; L6, L7 and L8 samples were collected at 34 dpi, L9 sample was collected at 41 dpi, L10 at 48 dpi, L11 and L12 at 54 dpi. M: Marker2000; EV: Empty vector (pV190). D, Relative expression level of *PDS* mRNA in the above indicated leaves determined by real-time qRT-PCR.

### The silencing effect of CGMMV-based vectors could be passaged

To verify whether the silencing vectors can be passaged, the sap of leaves with obvious photobleaching was used to rub-inoculate cotyledons and the L1 of melon plants. Photobleaching occurred on uninoculated leaves as early as 9 dpi and was photographed at 14 dpi (Fig. 6A). *PDS* expression levels were tested on L5 of passaged plants. We observed that *PDS* relative expression was reduced by 32%, 52%, 25% and 85% in pV190-PDS69, -PDS150, -PDS213 and -PDS300, respectively (Fig. 6B), confirming that the silencing effect of CGMMV-based vectors could be passaged.

**Figure 6:**
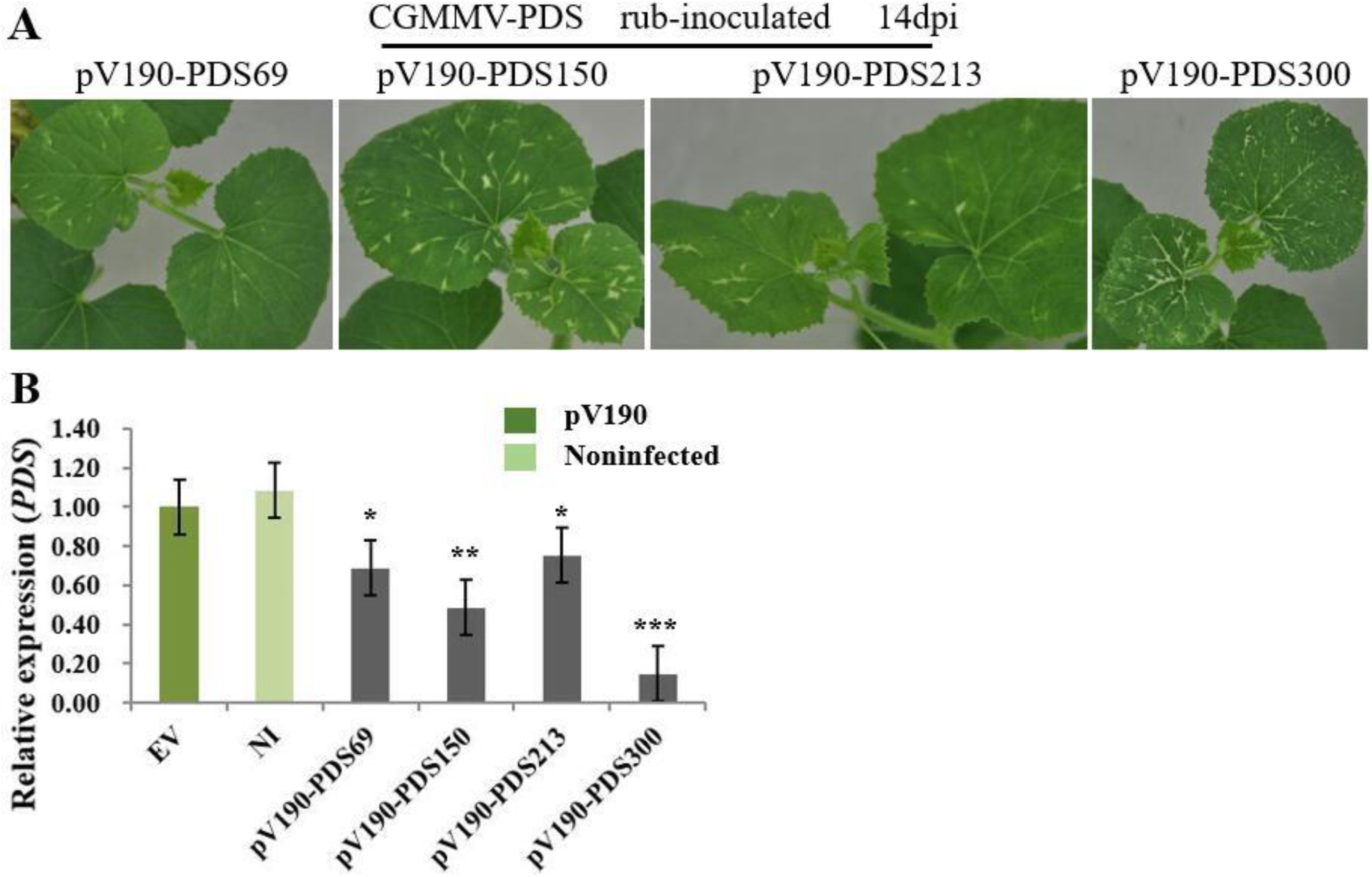
The silencing effect of pV190-PDS69, -PDS150, -PDS213 and -PDS300 could be passaged. A, Photobleaching caused by *PDS* silencing in systemic leaves of melon plants that were rub inoculated with sap from pV190-PDS69, -PDS150, -PDS213 and -PDS300-infected leaf tissue. The photobleaching phenotype was observed and photographed at 9dpi / 14dpi. B, Real-time qRT-PCR analysis of *PDS* expression in the 5^th^ leaf above the inoculated (L5) of noninfected (NI), pV190 empty vector (EV), and pV190-PDS69, -PDS150, -PDS213 and -PDS300-infected melon by mechanical inoculation.

### CGMMV-based VIGS in *N. benthamiana*

*N. benthamiana* is an important experimental host for CGMMV. We utilized two different lengths of *PDS* fragments which were amplified by selecting conserved regions of *PDS* gene sequences in *N. benthamiana* to test whether CGMMV is competent to induce gene silencing in *N. benthamiana* plants. At 14 dpi, a little weak photobleaching could be observed in the upper leaves of all plants inoculated with either pV190-NbPDS146 or pV190-NbPDS215 (Fig. S2A). Consistently, qRT-PCR results showed that the expression of *PDS* in pV190-NbPDS146- and pV190-NbPDS215-infected leaves was reduced by 60% and 34%, respectively, compared with the pV190 infected leaves (Fig. S2B).

## Discussion

In this study, we evaluated whether CGMMV could be used for constructing a viral vector to silence endogenous genes in cucurbit plants. A new CGMMV-based VIGS vector was developed through multiple attempts. Using this viral system, we successfully silenced *PDS* in cucurbits including watermelon, melon, cucumber and bottle gourd and in the model plant *N. benthamiana*. To our knowledge, this is the first time that CGMMV has been engineered as a VIGS vector, although it has been exploited for protein overexpression (Ooi et al., 2006; Teoh et al., 2009; Zheng et al., 2015; Jailani et al., 2017; Tran et al., 2019).

CGMMV has several characteristics that makes it a good candidate for VIGS vector in cucurbits. First, CGMMV can infect numerous species of cucurbits in natural conditions (Dombrovsky et al., 2017) and full-length infectious clones have been constructed successfully, which can systemically infect *N. benthamiana* and various cucurbit species including watermelon, melon, cucumber and bottle gourd and possesses high infection efficiency (*e.g.*, Liu et al., 2017). Second, CGMMV has a relatively small, positive single-strand RNA genome of 6,423 nt, which makes it easy to handle during either the process of preparing a VIGS construct or for inoculations.

During the process of modifying the CGMMV genome to produce a VIGS vector, we observed that the insertion sites of the gene fragment determined the viability, stability, insert size and silencing efficiency of the vector; our work showed that a duplicated copy of the 190-bp putative CGMMV CP SGP was essential for silencing. We first tried to place the foreign gene insertion site downstream of the viral CP gene. Results demonstrated that the insertion site between the CGMMV CP gene stop codon and the 3’ non-coding region was not suitable for constructing the VIGS vector. TMV is a member of the genus *Tobamovirus* and has successfully been developed as a VIGS vector by utilizing the strategy of subgenomic expression (Kumagai et al., 1995). Because CGMMV is also a member of the genus *Tobamovirus* and CGMMV infectious clone containing the green fluorescence protein (GFP) reporter gene has been successfully constructed, the GFP gene was located in between MP and CP (Zheng et al., 2015). Thus, we adopted a similar strategy to generate a CGMMV VIGS vector and explored different lengths of the duplicated region. When the modified CGMMV-based vector contained a duplicated copy of the 61-bp putative CGMMV CP SGP, the vector (pV61) lost its ability of systemic infection. When the modified CGMMV-based vector contained a duplicated copy of 92-bp or 112-bp putative CGMMV CP SGP, *PDS* silencing was not sufficiently robust (Table S3). In contrast, when the modified CGMMV-based vector contained a 190-bp duplicated copy of the putative CGMMV CP SGP, the *PDS* gene fragments could induce a robust silencing phenotype. These results suggest that it is necessary to create an additional fully competent subgenomic promoter to drive the transcription of the VIGS target sequence and for providing the vector with the ability to systemically infect plants (Mei et al., 2016).

Vectors containing duplicated sequences frequently suffer partial or complete loss of inserted sequences, particularly when the insert size is large (Avesani et al., 2007; Dickmeis et al., 2014). We tested the effect of the length and structure of inserts on silencing. Our results showed that the CGMMV vectors harboring the sense-oriented *PDS* gene sequence of 100-300 bp in length could effectively induce silencing in cucurbits, and efficiency was highest for largest fragment, the 300-bp *PDS* gene fragment. It is worth mentioning the effect on silencing of the cDNA insert length in a tobacco rattle virus (TRV)-based vector (Liu and Page, 2008). The better silencing phenotype could be produced when the cDNA insert length was between 200 bp and 1300 bp, whereas inserts shorter than 190 bp and longer than 1661 bp generated less siRNAs silencing less efficiently (Liu and Page, 2008). Not only the length of the insert affected silencing but also the structure of it has an impact on silencing. Expression of a hairpin-loop dsRNA structure could enhance the efficiency of VIGS (Lacomme et al., 2003). This seems to be true for our CGMMV-based VIGS vector. The silencing efficiency of a 69-bp hairpin-loop structure was between that of the 150-bp and 300-bp sense constructs, but its silence phenotype appeared earlier. A direct 60-bp inverted-repeat sequence of the target gene that could fold as dsRNA strongly enhanced VIGS from foxtail mosaic virus (FoMV) (Liu et al., 2016). However, in our work, a 60-bp inverted-repeat sequences of the *PDS* gene could not produce photobleaching, and CGMMV lost the ability of systemic infection (unpublished data). Therefore, these results suggest that the effect of length and structure of inserts on silencing varied with vectors from different viruses.

Furthermore, the stability of inserts in the pV190 vector were evaluated. The photobleaching phenotype was observed from the 3^rd^ to the 11^th^ leaves and *PDS* transcripts were reduced by about 80% and 20% in L4/L5 and L10/L11. About 70% of tested plants had a photobleaching phenotype which was stable and persisted for over two months. Stable photobleaching was also observed in plants mechanically inoculated with leaf sap prepared from L5/L6 of CGMMV-PDS inoculated plants. Here, it is worth mentioning that *PDS* transcript abundance could be reduced by CGMMV-PDS vectors on the 3^rd^ to 10^th^ leaves of the tested plants. However, the photobleaching phenotype and the *PDS* transcript levels were not uniform in these leaves, and could produce a gradient from bottom to top. It has been reported that the phenomenon could be due to instability of the *PDS* gene fragment. RT-PCR analyses on all of these leaves showed that deletion of the *PDS* insert was hardly detected in samples with the sense inserts. Hence, we reasoned that gene-silencing efficiency may be related to the accumulation of the foreign fragment-derived siRNAs (Molnar et al., 2010; Alvarado and Scholthof, 2012), but the specific mechanism of action is still unclear. In contrast, the full-length sequence of the insert in the pV190-PDS69 vector could not be detected in photobleached leaves. This phenomenon may be explained by a systemic silencing signal that can be actively transmitted over long distances through the phloem to induce *PDS* gene silencing in young leaves (Palauqui et al., 1997; Dunoyer et al., 2005; Molnar et al., 2010), but the exact molecular form of a mobile RNA signal in the phloem still needs further research.

In addition, we tested whether pV190 can be used to induce gene silencing in *N. benthamiana*. Results showed that pV190 could infect *N. benthamiana* leaves, the uninoculated systemic leaves developed very mild symptoms. The pV190 vectors, harboring 146-bp and 215-bp *PDS* fragments could trigger silencing, but the photobleaching phenotype was not striking. The photobleaching phenotype also varied in cucurbit plants, with the most obvious phenotype in bottle gourd. We reasoned that the viral vector has different fitness for different hosts. The accumulation of siRNAs is a crucial factor for silencing efficiency, while the host species also contain crucial factors including the *DCL, RDR* and *AGO* genes (Bouché et al., 2006) (Donaire et al., 2008; Zhang et al., 2012).

Apple latent spherical virus was first described as a vector for gene silencing in cucurbits (Igarashi et al., 2009), and few additional studies have reported its application for VIGS in cucurbit plants. A new TRSV vector was recently reported (Zhao et al., 2016). More recently, the TRV-VIGS system has been used in cucumber and oriental melon (Bu et al., 2019; Liao et al., 2019). Comparing ALSV, TRSV and TRV with CGMMV, the first three belong to multipartite virus families with a bipartite genome, while CGMMV is a monopartite virus (Ugaki et al., 1991), and therefore is easier to manipulate. The ALSV genome is expressed through polyprotein synthesis followed by proteolytic processing, which represents another layer of difficulty for high throughput functional genomics (Lu et al., 2003; Burch-Smith et al., 2004). A second major difference among the four cucurbit viral vectors is the inoculation method. *A. tumefaciens* infiltration is a simple, effective and convenient inoculation way (Ryu et al., 2004; Fu et al., 2005) and TRSV, TRV and CGMMV vectors are designed for agroinfiltration (Senthil-Kumar and Mysore, 2011; Zhao et al., 2016). Moreover, a new infection method with a special agroinfiltration solution used with the TRV VIGS system suitable for cucumber, provided rapidity, convenience and highly efficient gene silencing (Bu et al., 2019). Host range is another major difference. TRSV cDNA clones are not infectious in watermelon or pumpkin (Zhao et al., 2016). Although TRV has been widely used as VIGS vector since it has a wide host range (Ratcliff et al., 2001), its application for VIGS on cucurbits except cucumber and oriental melon have not been reported (Bu et al., 2019; Liao et al., 2019). Both ALSV and CGMMV vectors can be successfully used on common cucurbit plants such as watermelon, melon, cucumber and bottle gourd (Igarashi et al., 2009). At last, the CGMMV-based VIGS vector pV190 can produce very mild viral symptoms in upper leaves of inoculated plants and is highly infectious. Silencing phenotypes caused by pV190-based VIGS vectors were stable and could persist for at least one month.

CGMMV has a broad host range including 29 species, of which at least 16 belong to Cucurbitaceae (Dombrovsky et al., 2017). Thus, the pV190 VIGS vector should have the potential for VIGS in many other plants although we only evaluated its application in *N. benthamiana*, watermelon, melon, cucumber and bottle gourd. Taken together, the CGMMV-based silencing system could be applied as a powerful biotechnological tool with a great potential for studying functional genomics in cucurbits. Future study will focus on obtaining insights into the molecular mechanism underlying the difference in silencing efficiency between different plants. In addition, the vector could serve as a basis to control devastating viral pathogens or carry out genetic engineering and molecular breeding.

## Accession Numbers

Sequence data from this article can be found in the GenBank or Cucurbit Genomics Database (http://cucurbitgenomics.org/) under the following accession numbers: CGMMV (KC851866); *Nicotiana benthamiana PDS* (EU165355); *Cucumis sativus PDS* (XM_011654729); *Cucumis melo PDS* (NM_001297530); *Citrullus lanatu PDS* (Cla010898; ClCG07G015130); *Lagenaria siceraria PDS* (Lsi07G003470).

## Supplemental Data

**Supplemental Figure S1:** The sequence similarities of 114-, 150-, 213- and 300-bp *PDS* gene fragments in four cucurbit species. A, B, C and D correspond to *PDS* fragments of 114-, 150-, 213- and 300-bp, respectively, in the four cucurbit species. HG, HUG, XG and TG represented cucumber, bottle gourd, watermelon and melon, respectively.

**Supplemental Figure S2:** Silencing efficiency of different length inserts (*PDS*) using the pV190 VIGS vector in *N. benthamiana*. Fragments of 146 bp, 215 bp were separately cloned into pV190 VIGS vector. A, The silencing phenotypes were observed at 14dpi. B, The relative expression level of *PDS* mRNA determined by real-time qRT-PCR.

**Supplemental Table S1:** Primers used in this study.

**Supplemental Table S2:** The infection analysis of pV1a23 (insertion sites behind the viral CP gene) vector in watermelon.

**Supplemental Table S3:** The infection analysis of modified CGMMV-based vector contains a duplicated copy of the 61-, 92-, 112- and 190-bp putative CGMMV CP SGP.

## ACKNOWLEDGMENTS

We thank Yule Liu (Tsinghua University, Beijing) for thoughtful advice on designing VIGS vectors. We also thank Tao Zhou (China Agricultural University, Beijing) and Zhangjun Fei (Boyce Thompson Institute, USA) revised this manuscript.

## Notes

**Funding information:** This research was supported by the National Natural Science Foundation of China (31571247), the grants from the earmarked fund for the China Agriculture Research System (CARS-26-13), and the Agricultural Science and Technology Innovation Program (ASTIP), Chinese Academy of Agricultural Sciences (CAAS-ASTIP-2018-ZFRI-08).

